# Laboratory study of *Fritillaria* lifecycle reveals larvacean commonalities and key morphogenetic events leading to genus-specific anatomy

**DOI:** 10.1101/2022.05.25.493221

**Authors:** Simon Henriet, Anne Aasjord, Daniel Chourrout

**Affiliations:** Sars International Centre for Marine Molecular Biology, University of Bergen, 5006 Bergen, Norway

## Abstract

A fascinating variety of adult body plans can be found in the Tunicates, the closest existing relatives of vertebrates. A distinctive feature of the larvacean class of pelagic tunicates is the presence of a highly specialized surface epithelium that produces a cellulose test, the “larvacean house”. While substantial differences exist between the anatomy of larvacean families, most of the ontogeny is derived from the observations of a single genus, *Oikopleura*. We present the first study of *Fritillaria* development based on the observation of individuals reproduced in the laboratory. Like the other small epipelagic species *Oikopleura dioica*, the larvae of *Fritillaria borealis* grow rapidly in the laboratory, and they acquire the adult form within a day. We could show that major morphological differences exhibited by *Fritillaria* and *Oikopleura* adults originate from a key developmental stage during larval organogenesis. Here, the surface epithelium progressively retracts from the posterior digestive organs of *Fritillaria* larvae, and it establishes house-producing territories around the pharynx. Our results show that the divergence between larvacean genera was associated with a profound rearrangement of the mechanisms controlling the differentiation of the larval ectoderm.

---

Larvaceans, also known as appendicularians, are a class of tunicates characterized by the production of a unique filter-feeding apparatus, nicknamed the “larvacean house”. The house is a complex structure secreted by the animal, which consists in multiple rooms whose function is to size-select food particles (usually microalgae and bacteria) and to channel the water flow in or out from the digestive tract^1^. Discarded houses will sink and aggregate organic matter, thus fast-growing larvacean populations during algal bloom will contribute to the transfer of significant amounts of carbon to the sea floor^2^. While the mechanisms that drive the assembly of the cellulose and protein mesh structure remain largely unknown, some of the anatomical and molecular basis for house synthesis have been uncovered recently - mostly by studying the species *Oikopleura dioica*. Results obtained with this species have revealed important evolutionary prerequisites such as the acquisition of bacterial cellulose synthase genes, the emergence of a novel family of proteins - the oikosins^3^; and the deployment of novel genetic networks involved in the differentiation of cell territories within the house-producing organ, also known as the oikoplastic epithelium^4,5^.

Zoological records have permitted to establish three larvacean families, the Oikopleuridae, the Kowalevskiidae, and the Fritillariidae. The diversity of these organisms has been studied mostly by examining specimens freshly caught in the open ocean and brought to the surface on exploration vessels, which usually do not offer suitable conditions for reproducing captured specimens and studying lifecycles. Thus, apart from anatomical descriptions that have often been conducted with rudimentary microscopy techniques, the biology of many larvaceans remains obscure. The few exceptions are found in the Oikopleuridae, and they include *Oikopleura dioica*, a coastal species that can be raised in the laboratory^6,7^. ROVs may grant better access to larvaceans biology *in situ*, and they have been deployed for studying giant oikopleurids off the Northwest American coast^8^. While these recent advances have provided precious insights about the evolution and development of the taxon^9^, there is a contrasting lack of knowledge for other families. Modern descriptions of the Kowalevskiidae are scarce^10,11^ and recent studies of fritillarids have focused on the few features that can be accessed directly, like gut anatomy and animal behavior during house production^12–14^. Due to the lack of comparative information, we may under-appreciate the variety of biological processes present in larvaceans, which in turn compromises our understanding of their evolutionary history at the base of the vertebrate lineage^15^.

The genus *Fritillaria* contains interesting candidate species to address some of the knowledge gaps. At the ecological level, *Fritillaria borealis* shares similarities with *O. dioica*, such as geographical and seasonal distribution, and presence in coastal surface waters. At the genetic level, *F. borealis* and *O. dioica* are characterized by small and fast-evolving genomes, a likely result of large effective population size and other mechanisms driving chromosome reduction^4,16^. There is a major difference at the level of reproduction since *O. dioica* is the only non-hermaphroditic tunicate. Like all larvaceans, the general anatomy is relatively simple and consists in a trunk with feeding, reproductive and sensory organs, followed post-anally by a long tail used by the animal for locomotion and to generate water flow in the house (**Fig1**). At the end of larval development, the trunk of *Oikopleura* assumes a round compact shape. It consists in an anterior cavity that includes all digestive organs, covered with the oikoplastic epithelium^12,17^, and followed posteriorly by a cavity with reproductive organs whose size and complexity will increase dramatically during sexual maturation^18^. At a comparable stage in *Fritillaria*, the trunk has a very different, elongated shape that can be divided in three segments along the antero-posterior (AP) axis^19^. The anterior-most segment consists in a cavity covered with oikoplastic epithelium, that contains the mouth, the cerebral ganglion, and the pharynx. It is connected by the oesophagus - which consists in a single layer of small cells, to a central segment which is a simplified gut composed of few large cells^20^. The posterior-most segment corresponds to the gonad sac, which grows inside a cuticular extension of the trunk. There is no epidermis visible in segments posterior to the pharynx, and the only barrier to the outer environment is a thin cuticular layer without visible cellular structures. *Fritillaria* and *Oikopleura* also exhibit fundamental differences during house production, which could influence their respective contribution to “marine snow” aggregates. In *Oikopleura*, the house expands only once after its synthesis, it surrounds the whole animal, and it is frequently discarded (usually following clogging or stress)^21,22^. In *Fritillaria*, the house expands on the anterior side only. The animal is capable to deflate and re-inflate the house at regular time intervals, which is believed to facilitate filter function by chasing clogged particles^13^.

**Figure 1:**
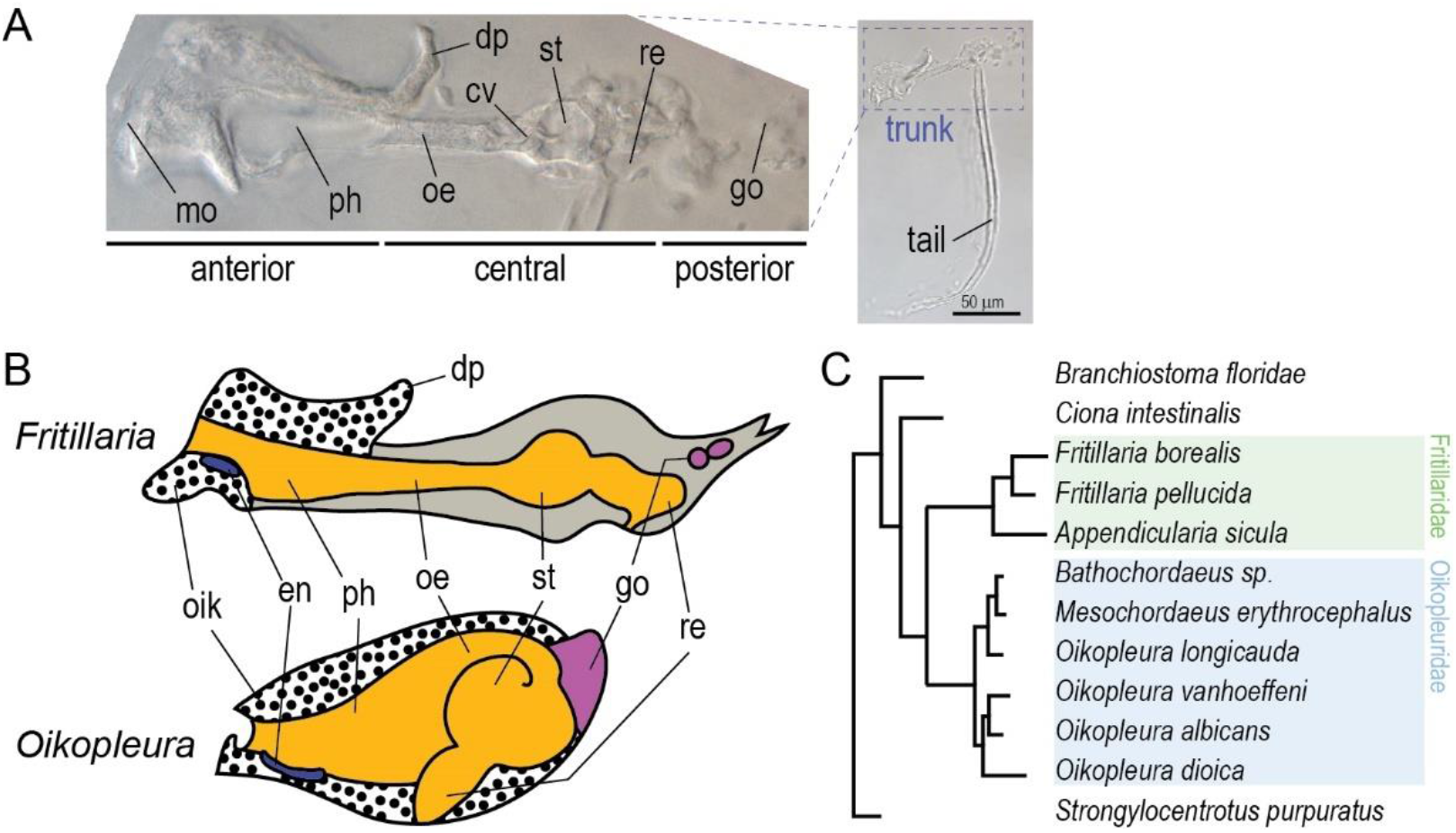
Anatomy of the trunk in two larvaceans families. **A**) Young juvenile *F. borealis* (2 days post fertilization (dpf)), with detail of the trunk organs. **B**) Schematic representation of the trunk organs in *Fritillaria* and *Oikopleura*. **C**) Phylogeny of larvaceans adapted from Naville *et al*^16^. en, endostyle; cv, cardiac valve; go, gonad; oe, oesophagus; oik, oikoplastic epithelium; ph, pharynx; st, stomach; re, rectum.

Taking advantage of a laboratory culture maintained over several months, we have gained better insight on some of the known characteristics of *Fritillaria*. This includes the peculiar trunk anatomy - previously described in adults only^20^, the development of the female gonad - so far documented with low-resolution microscopy techniques^23^, and the short lifecycle whose duration was inferred from plankton tows^24^. By using confocal microscopy to gain detailed access to the larval anatomy, we could reveal common features also observed in *O. dioica*, and key differences during organogenesis. Our results show that the distinctive anatomy of *Fritillaria* is determined by events that take place during the development of the trunk in hatched larvae. The mechanisms appear to involve the differentiation of the trunk in three distinct regions that are analogous to the segments observed in adult animals, prior to the progressive dissociation of the larval ectoderm from the posterior side of the trunk. These morphogenetic processes are not observed in *O. dioica*, and they can clearly account for the organization of digestive organs and house-producing epithelium in the adult form of *F. borealis*.

## 1 Reproduction of *F. borealis* in the laboratory

We collected *F. borealis* from surface tows at three locations in the fjords around Bergen in western Norway. Sea water samples were transported to a laboratory facility and grown under constant paddle agitation at 12°C^6^. With naked eyes, it is possible to identify few animals with developing gonads already immediately after collection, and this is often the case during spring and autumn when population densities are higher^25^. However, water samples collected during these seasons also contain numerous larvae and small juveniles which become visible after two days of growth in the lab. These young animals make a larger contribution to the initial increase of individuals in culture, and their presence is usually a favorable prognostic for maintaining laboratory populations over a longer term.

Two forms of parasites were found associated with wild specimens (**Fig2A**). The first form is easily seen on mature animals and consists in a row of bead-shaped cells that replace the gonads in the posterior trunk. These cells easily detach from the animal (for example upon transfer to an observation container) and when isolated, they rapidly divide into ciliated swimming forms. This parasite is most likely a Syndiniale of the genus *Sphaeripara*, ectoparasitic dinoflagellates previously described on *F. pellucida* specimens^26^, and whose interaction with *F. borealis* was shown indirectly with metagenomics^27^. The second form is visible only under microscopy and consists in single, round cells attached to the trunk. It probably corresponds to dinoflagellates ectoparasites of the genus *Oodinium*^28^, which we also found on *O. dioica* specimens collected in the same areas.

**Figure 2:**
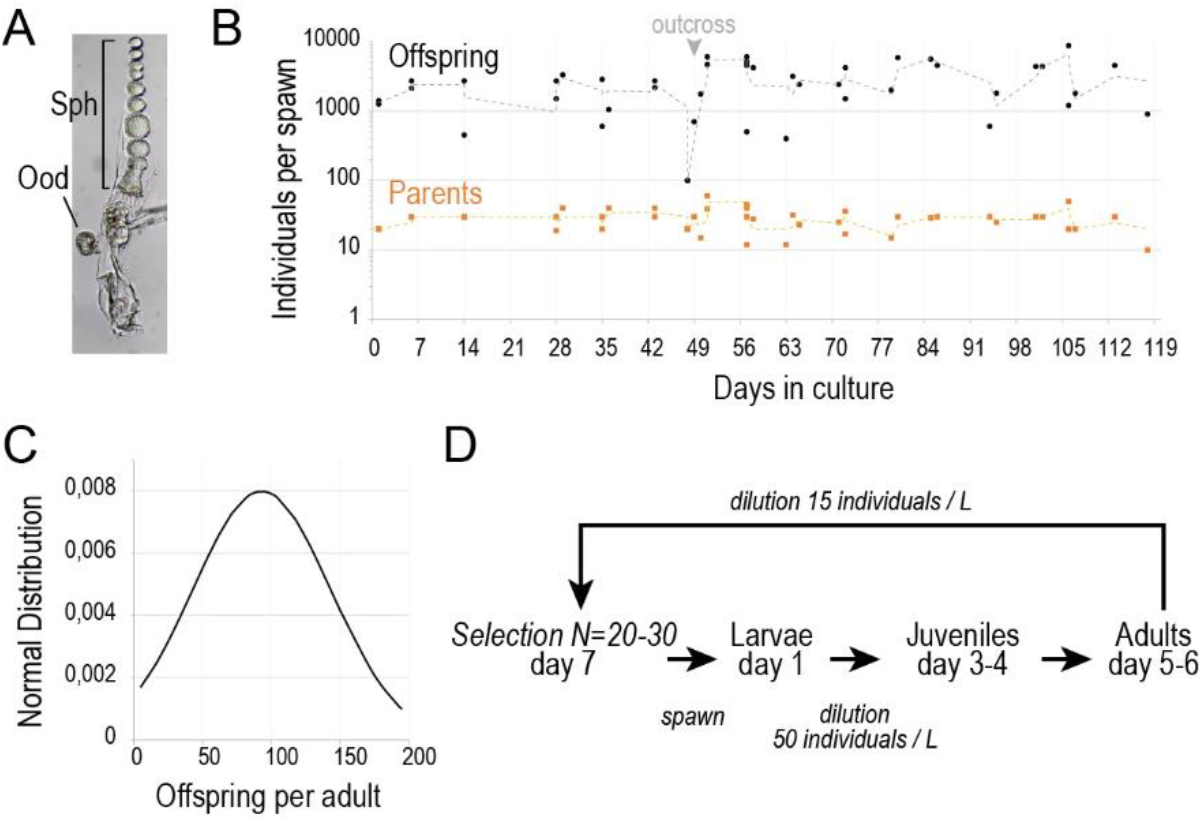
Laboratory culture of *F. borealis*. **A**) Specimen collected from the wild, with parasites. **B**) Recordings of spawns conducted with laboratory animals during a 17-weeks period. The number of adults and offspring produced (counted from 1 dpf samples) has been plotted for each spawn. Curves show 2 periods moving averages for each series. Laboratory populations were bred excepted at week 7, where spawns were established between two populations. **C**) Average fecundity of adults determined from the offspring counted after laboratory spawns. **D**) Schematic of the culture procedure showing the seven days lifecycle and animal handling (italic font).

Several attempts have been made to reproduce and maintain the animals over multiple generations. Parasitism was a leading cause of failure, prompting us to systematically screen for parasite-free animals during the isolation from field sample. Ciliates can have non-antagonistic relationships with *O. dioica*^29^ but in the case of *F. borealis*, we found them more likely to represent a nuisance and we addressed their proliferation by using sterile growth media for the algae cultures used as food source. Feeding *F. borealis* with the pico-algae *Micromonas, Phaeocystis* and *Synechococcus* gave the best results, while a diet of larger cells such as the one optimized for *O. dioica*^6^ led to poor animal growth, possibly due to inadequate size of food particles. Measures to prevent parasite and ciliate infestation, together with the appropriate diet, have proven decisive to maintain fecundity and to extend the laboratory culture over several weeks (**Fig2B**).

## 2 Germline development and fertilization

Larvaceans are semelparous animals that reproduce by external fertilization. In the laboratory, mature founders for the next generation are selected at the end of the seven days lifecycle after examining body size and gonad morphology (**Fig2D**). Two days after fertilization, gonads appear as undifferentiated rudiments surrounded by transparent cuticle (**Fig3A**). The composition of the cuticle is unknown and looks similar to the tail fin. After four days of growth, it is possible to differentiate the testis, located in an apical position, and the spherical ovary placed close to the gut. The cuticular extension is completely occupied by gonads at this stage, apart from a pair of distal, horn-shaped appendages whose supposed function is to provide anchor points for the house^13^. Based on DAPI staining and previous observations on *O. dioica*, we identified nurse nuclei with high amount of peripheral chromatin^30^ (**Fig3B**). Their size is initially large in immature gonads, and they lie within an extensive actin network reminiscent of the *O. dioica* coenocyst^18^. Like *F. pellucida*^23^, female germline nuclei are absent from this network and at later stages, oocytes indeed form an outer layer that surrounds the cavity where nurse nuclei are located (**Fig3C**). It suggests that at early stage, female germ cells could be already present in the outer layer of the ovary, possibly among numerous somatic cells. The alternative would be that female germ cells that are initially located in the coenocyst, migrate towards the periphery to complete their development into oocytes. This idea appears inconsistent with the presence at late stages, of a thick extracellular shell that separates the growing oocytes and the nurse nuclei. Nurse nuclei are dramatically reduced before gamete release, but they remain visible as DAPI-positive material between oocytes (**Fig3D**). At this stage, oocytes exhibit a prominent germinal vesicle with four clearly distinguishable chromosomes, a number consistent with the *F. pellucida* karyotype^31^ (**Fig3E**). Immediately before spawn, the ovary ruptures and oocytes invade the gonad cavity. Gametes are subsequently released in the seawater through an apical opening of the gonad, and during this process the eggs are being forced through the remnants of the testis sac. After oocyte release, DAPI-positive spots can be observed at the surface. Even though we cannot completely exclude that these may correspond to follicular cells, which are frequently encountered in ascidians, their small size and variable number (between 5 and 12 per egg) also suggest they could represent leftover of nurse cell nuclei stuck to the oocyte.

**Figure 3:**
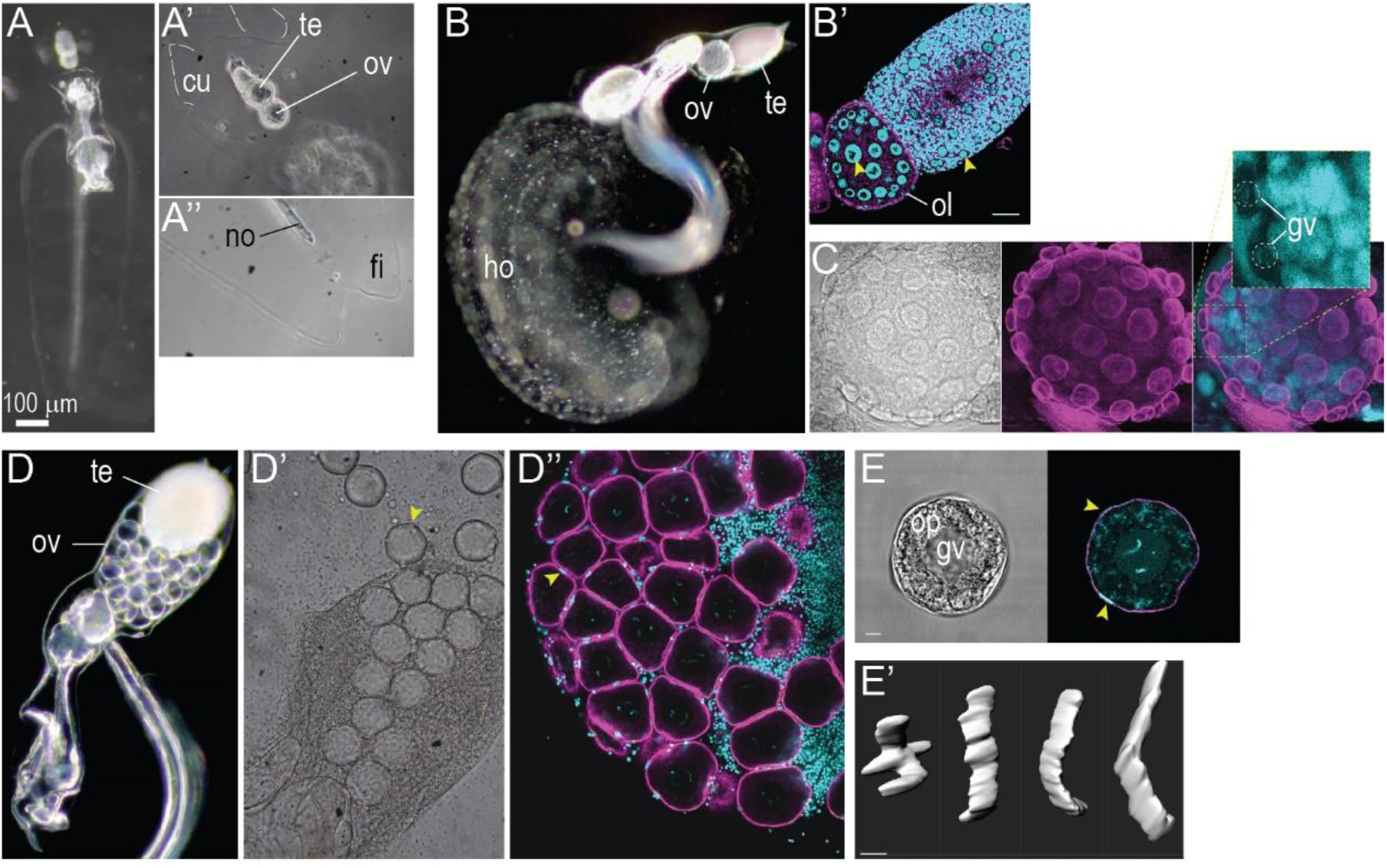
Gonad development in *F. borealis*. **A**) Specimen observed early during the lifecycle (2-3 dpf), with detail of the cuticle around the gonad (**A’**) and at the tip of the tail (**A’’**). **B**) Specimen observed at an advanced stage of development (5 dpf). Yellow arrowheads indicate nurse nuclei in the ovary and testis (**B’’**). **C**) Ovary observed late during the lifecycle (6-7 dpf), showing growing oocytes at the periphery of the ovary. **D**) Specimen close to spawn with mature gametes (7 dpf), and observation of the ruptured gonad (**D’**, **D’’**). **E**) Oocyte observed immediately after release and 3D reconstruction of the female chromosomes (**E’**). Cyan, DAPI; magenta, phalloidin-AF488; cu, cuticle; fi, tail fin; gv, germ vesicle; ho, house; no, notochord; ol, outer layer of the ovary; op, ooplasm; ov, ovary.

In ascidians, fertilization is controlled by a non-self/self recognition mechanism that operates at the surface of the egg^32^. Many species are self-sterile, and outcrossing is likely predominant in natural conditions^33^. Self-fertilization of *F. borealis* was never observed in our hands (N = 10), whereas offspring could be recovered from crosses between single individuals >60% of the time (N = 10). Development speed during early embryogenesis is comparable to *O. dioica*, with rapid blastomere divisions after fertilization (every 30 minutes at 15°C) and hatching of swimming larvae between 4-6h after fertilization.

## 3 Anatomy of the house-producing epithelium

EM sections have offered valuable insights into the anatomy of the different larvacean families^11,20,34^. Techniques are generally well-suited for whole-mount observations of small animals, and the simple preparation of specimens represent safer procedures when samples are in limited amount. Whereas EM permitted to gain exquisite knowledge about the gut anatomy^11,20,35^, it brought comparatively little new information about the organization of the house-producing epithelium. For this matter in particular, techniques based on fluorescent probes capable to reveal filamentous structures present in larvacean houses appear more appropriate.

We used a combination of DAPI and phalloidin staining to reveal individual cells in the trunk of *F. borealis*. This approach permitted to easily identify feeding organs and the nervous system (which we also stained with immunofluorescence) (**Fig4A**). However, using this approach only proved unconclusive to directly attribute house-producing function to specific epithelial cells. Unlike oikopleurids, whose epithelium shows well-delimited fields of cells with characteristic shape and size^36^, most of the *F. borealis* surface cells are homogenous in shape excepted for a pair of dorsal rows of cells located centrally on the pharyngeal trunk epithelium. Moreover, the flattened and curved shape of the trunk proved to be problematic for distinguishing superficial cells from the tissues located underneath. Confocal scanning proved useful for address the latter issue, while we used a fluorescent Carbohydrate-Binding Module (CBM) to stain cellulose and identify cells related to house production. Cellulose fibrils originate from different locations on the surface of the anterior trunk. Many cellulose-secreting cells are located in a “horseshoe-shaped” projection of the epidermis that extends laterally and dorsally over the oesophagus (**Fig4A, B**). These cells are significantly larger, suggesting that Fritillarids and Oikopleurids may share conserved mechanisms to regulate cell size in specific territories of the epithelium^37^. Fields of house-producing cells were also found dorsally and ventrally around the mouth in the anterior part of the epidermis. In *F. borealis*, we could not distinguish cellulose staining on the dorsal rows of cells. However, their location strongly suggest they could contribute to the formation of specific structures in the house, such as a pair of dorsal chambers easily seen in animals whose house has not yet been inflated (**Fig4A’**). The fluorescent CBM did not stain such chambers and was not reactive either to the gonad and tail cuticle (**FigS**). Outside the spiracle opening and the anterior house-producing fields, the ventral side appears extremely simplified and most of its surface is occupied by two giant cells whose function remains unclear (**Fig4B**).

**Figure 4:**
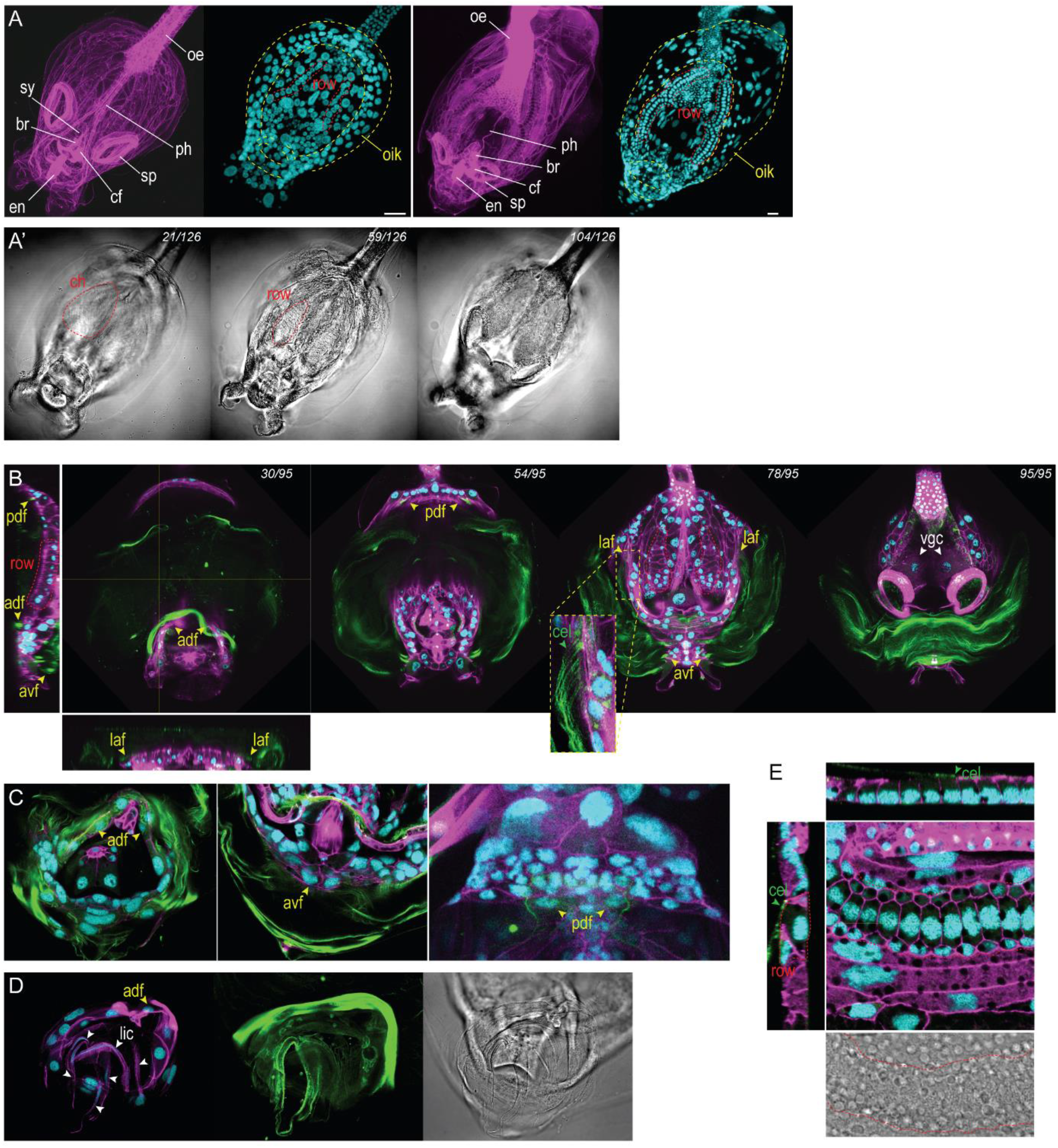
House-producing cells in post-larval stages of Fritillarids. **A**) Maximum intensity projection of the anterior trunk of *F. borealis* (left) and *F. haplostoma* (right) specimens, showing the main internal organs (left) and fields of epidermal cells (right). Scale bar, 20 microns. Optical sections through the anterior trunk of the *F. borealis* specimen (**A’**) showing chambers of the house, surface cells located underneath on the dorsal side, and the ventral side with the spiracle openings. **B**) Optical sections of the anterior trunk with fluorescence staining of the house. Left and bottom panels show orthogonal projections of the Z-stack. Yellow arrowheads indicate fields of house-producing cells, green arrowheads show projections of cellulose fibrils. **C**) Conserved fields of house-producing cells in *F. haplostoma*. **D)** The lip organ in *F. haplostoma*. Panels from left to right show: composite DAPI and AF-488 fluorescence, GFP fluorescence, and bright field illumination. **E)** Dorsal epithelial row of cellulose-secreting cells in *F. haplostoma*. Left and upper panels show orthogonal projections of the Z-stack. Bright field illumination (bottom panel) shows the opaque vesicles present in epithelial cells. Cyan, DAPI; green, GFP-CBM; magenta, phalloidin-AF488; adf, anterior dorsal field; avf, anterior ventral field; br, brain; ce, cellulose fibrils; cf, ciliary funnel; ch, house chamber; en, endostyle; li, lip; laf, lateral field; lic, lip cell; oe, oesophagus; oik, oikoplastic tissue; pa, palps; pdf, posterior dorsal field; ph, pharynx; row, dorsal row of epidermal cells; sp, spiracle; sy, statocyst; vgc, ventral giant cell.

We conducted the same analysis on individuals from another species, identified as *Fritillaria haplostoma* on the basis of based on tail and gonad morphology (**FigS**)^19^. House-producing cell fields were found at conserved positions on the pharyngeal trunk (**Fig4C**). The larger body size allowed to better observe cellulose fibrils, which permitted to confirm the role of the dorsal rows of cells in house secretion. We also observed fundamental differences with the anatomy of *F. borealis*, possibly related to different house structure and feeding behavior. Thus, the pair of lateral palps flanking the *F. borealis* mouth is absent in *F. haplostoma*. Instead, we observed a delicate arrangement of cellulose-rich, flattened cells forming a protruding structure around the mouth opening which could corresponds to the “lip” mentioned in previous descriptions^19^ (**Fig4D**). The position of these organs indicates a function during feeding, possibly for processing specialized diet. We could not directly compare morphology and number of house-producing cells in the two species. This is partly due to the orientation of the specimens, and to the unknown developmental stage of *F. haplostoma* specimens that were obtained from field collection. However, we could clearly observe that the dorsal rows of cells of *F. haplostoma* assumed a more complex arrangement, which consists in two rows of small cells flanking a central row of larger, rectangular cells (**Fig4E**). Unlike *F. borealis*, we also observed numerous, large opaque vesicles that are distributed uniformly across the pharyngal trunk epithelium of *F. haplostoma*. We suppose these organelles could participate to epithelium function, like secretion of house components. Alternatively, the opaque content could indicate a protective function against UV radiation.

## 4 Morphogenesis of the trunk epithelium

While all larvaceans can produce filter-feeding houses, it remains unknown if the development of the oikoplastic epithelium is controlled by common mechanisms in different larvacean families. Developmental processes have been documented mainly in *O. dioica*, showing that epithelium morphogenesis occurs early during development and several hours before the first house is synthesized^5^. While such timing could be conserved during the development of *F. borealis*, the presence of larvacean family-specific oikoplastic fields in adult animals strongly suggest that distinct developmental mechanisms also exist. We decided to gain better insight by examining the development of the epithelium in *F. borealis* larvae. Direct observation of developing embryos proved challenging, as the development of *in vitro* fertilized oocytes frequently aborted when animals were left in vessels without water flow. This issue prevented us to document morphogenetic events taking place during blastula and gastrula stages with accuracy (**FigS**). However, when fertilization took place in spawn vessels under agitation, we could efficiently recover hatched embryos and swimming larvae. The developmental progression of these animals could be determined *a posteriori* under microscope, by examining key features such as cell number, notochord growth and epithelium patterning.

Trunk and tail primordia can be recognized before hatching, when the embryo assumes a coiled shape with two lobes that resembles the tailbud stage of *O. dioica*^9^ (**Fig5A**). In these animals, the notochord is already visible and consists in 12 to 17 cuboid cells stacked along the AP axis. In the next stage observed, the tail and trunk are more differentiated and the notochord cells have completed their divisions, reaching a final number of 20 cells (including a round-shaped terminal cell in the tip of the tail), equal to the *O. dioica* notochord^9^ (**Fig5B**). The tail anatomy features characteristic cell types also observed in *O. dioica*, such as broad muscle cells flanking the notochord, and a small group of cells located on one side of the tail surface that will later give rise to the caudal ganglion. In contrast, tissues of the trunk remain hard to characterize, apart from columnar cells that will later give rise to the epithelium.

**Figure 5:**
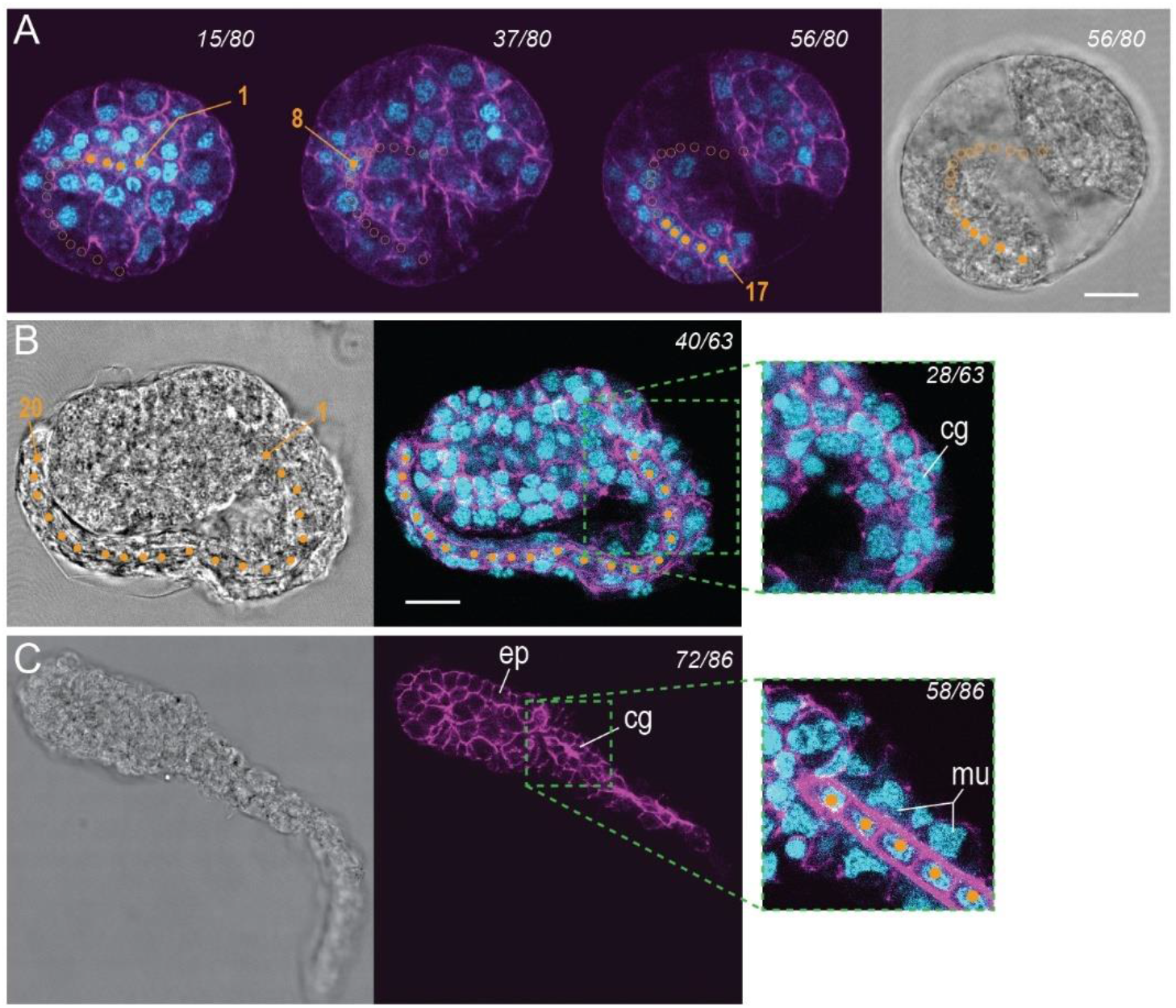
Early stages of notochord and tail development in *F. borealis*. **A**) Optical sections showing an embryo at the tailbud stage, with notochord cells stacked along the AP axis (orange dots). Full dots represent visible cells, open dots represent cells visible in other sections. **B**) Embryo at the end of notochord cell divisions. **C**) Embryo immediately after hatching. cg, caudal ganglion; ep, epidermis; mu, muscle cells. Scale bar, 10 microns.

The larvae of *F. borealis* and *O. dioica* look very similar immediately after hatching (**Fig5C**). However, their morphology differs dramatically a few hours later, at a developmental stage that also coincides with elevated swimming activity. In *F. borealis*, the trunk differentiates in two lobes that will respectively form the pharynx and the gut. These lobes are separated by a central constriction that corresponds to the developing oesophagus (**Fig6A**). The segmentation of the larval trunk and its extension along the AP axis thus appears as first morphogenetic processes accounting for the characteristic adult morphology (**Fig1**). In contrast, *O. dioica* larvae observed at a similar stage have a compact trunk, where the presumptive gut folds back towards the anterior and develops ventrally under the oesophagus (**FigS**). We observed the intriguing presence of surface fibrils running between the anterior and posterior lobes on the dorsal side of the *F. borealis* larva, which suggests that epithelial cells are already carrying out some functions. Supporting this idea, distinctive cell arrangements previously seen in house-producing animals (**Fig4B**) can be recognized on the dorsal epithelium of the anterior lobe (**Fig6A**) and on the ventral epithelium located under the mouth opening. It is possible that the surface fibrils secreted by the larval ectoderm could contribute to the “cuticular layer”^38^ that has been observed around the animal.

**Figure 6:**
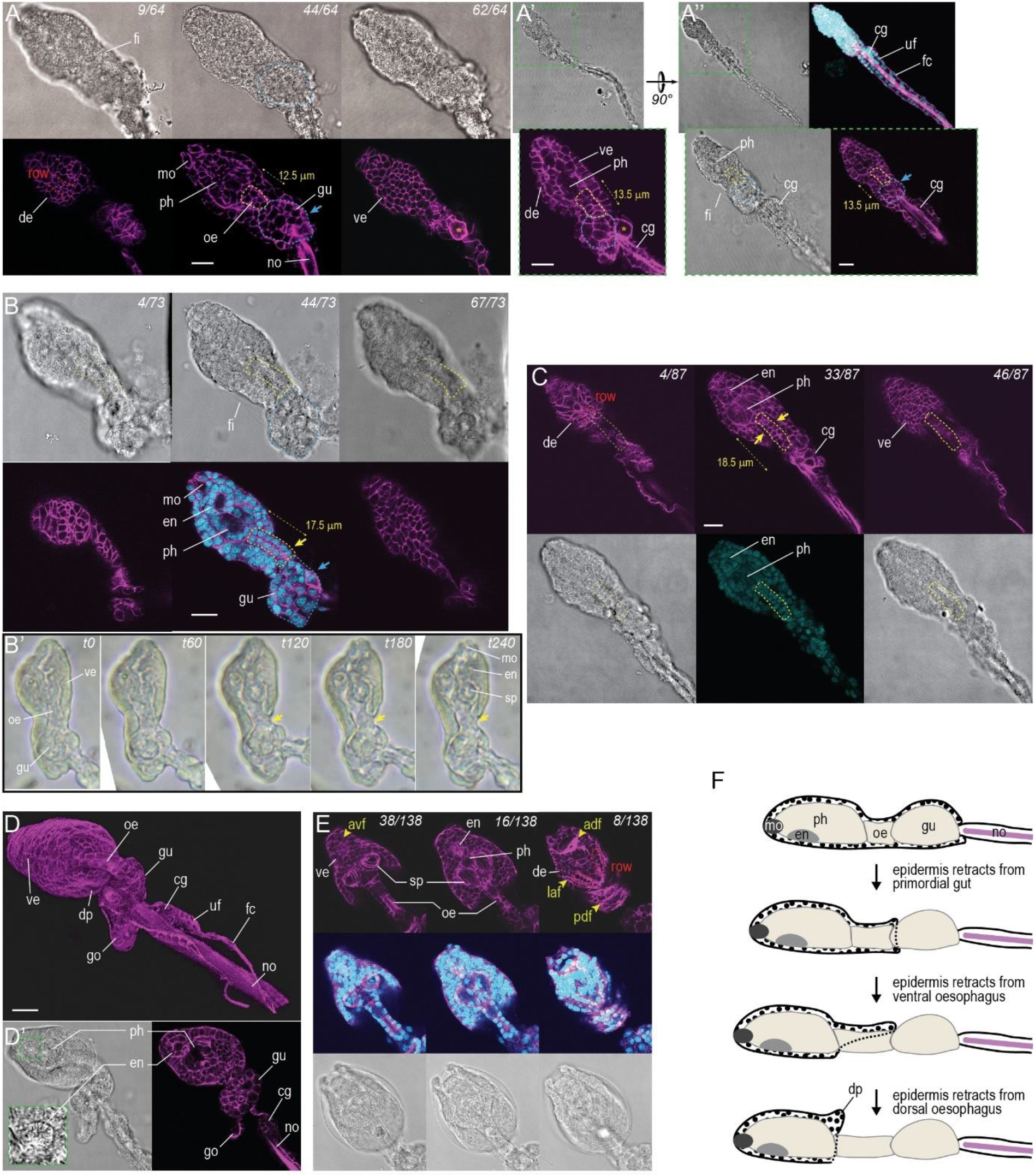
Morphogenesis of the trunk in *F. borealis*. **A**) Optical sections on the dorsoventral axis, side view (**A’**) and dorsal view (**A”**) of hatched larvae observed during the first step of epidermis retraction. Note the thickened basal lamina stained with phalloidin (magenta), used to outline the position of the epidermal layer. Yellow and blue dotted lines mark the position of developing oesophagus and gut, respectively. Blue arrows show area of the gut where epidermis has retracted. The star marks a large cell located on the ventral side which is used to orient the larva. **B**) Optical sections and movie frame shots (**B’**) of larvae observed during the second step of epidermis retraction. The epidermal layer is completely absent around the gut, and it is also retracting from the ventral side of the oesophagus (yellow arrows). **C**) Optical sections of a larva observed at the end of the second step of epidermis retraction. The epidermal layer is dissociated from the surface of the oesophagus. **D**) 3D reconstruction and optical section (**D’**) of a larva during the third step of epidermal retraction. The epidermis has formed a projection over the dorsal side of the oesophagus. The gonad primordium is now visible in the posterior trunk, lined by a cuticular material similar to the tail fin. **E**) Juvenile with a pre-house. In this animal collected within 24hpf, it is possible to recognize the major anterior organs and house-producing fields of cells on the trunk epidermis. adf, anterior dorsal field; avf, anterior ventral field; laf, lateral field. **F**) Schematic description of the epidermis retraction during larval development. cg, caudal ganglion; de, dorsal epidermis; dp, dorsal projection; en, endostyle; fi, surface fibers; fc, fin cell; go, gonad; gu, gut; mo, mouth; no, notochord; oe, oesophagus; ph, pharynx; sp, spiracle; uf, upper fin cell; ve, ventral epidermis. Scale bar, 10 microns

In *F. borealis* and in *O. dioica* as well, the larval epithelium initially covers all inner tissues of the trunk, excepted the mouth (**Fig5C**, **FigS**). Throughout larval development, the trunk of *O. dioica* will remain covered by ectoderm, excepted at its posterior end where germline precursors will later give rise to the gonad (**Fig1**). By examining hatched larvae collected at successive developmental stages, we could document that in *F. borealis* the trunk epithelium progressively retracts from the posterior end towards the anterior, leaving exposed the inner tissues. We distinguished three steps during this process. The first step appears to coincide with oesophagus elongation along the AP axis. Retraction of the epithelium could be driven by the proliferation of oesophagal cells (**Fig6A, B, C)** and it concludes by a complete absence of outer cell layer around the gut primordia. The second step was observed after organs of the anterior trunk, such as like the endostyle and the pharynx, have acquired their characteristic morphology. Here, the ectoderm dissociates progressively from the oesophagus and leaves the ventral side first (**Fig6B**). A multi-layered mass of cells is still visible on the dorsal side, suggesting that the retraction could be driven by the collective migration of ectodermal cells. At the end of the second step, the oesophagus has lost all direct contacts with the epithelium, but it remains surrounded by epithelial tissue (**Fig6C**). During the third step the epithelium forms the horseshoe-shaped, dorsal projection over the oesophagus (**Fig6D**), where house producing cells have been evidenced. On the ventral side, the number of epithelial cells is dramatically reduced after the spiracles have appeared, suggesting that some epithelial cells could contribute to spiracle formation (**Fig6E)**.

## Discussion and conclusions

Tunicates are a fascinating group of animals with considerable diversity in size, feeding strategy, reproduction, and other fundamental life traits. However, biological studies have been more sustained on sessile tunicates (represented by solitary and colonial ascidians) than on pelagic tunicates, even though the latter are increasingly recognized as key components of the oceanic ecosystems^39^. Some pelagic tunicates are difficult to access and as a matter of fact, the abundant and ubiquitous *O. dioica* rapidly became one the few species to be established in a continuous laboratory culture^6^. Studying the *O. dioica* genome revealed many surprising features and illuminated some of the molecular bases underlying the biology of this species. However, it is by employing a comparative approach based on a variety of larvacean genomes^16^ that we could uncover general mechanisms explaining the evolutionary trajectory of larvaceans. Under this perspective, bringing *F. borealis* to the laboratory represents a decisive step for testing the conservation of biological processes between larvaceans families. By significantly improving the level of detail of the *Fritillaria* anatomy, our study provides first results that should be useful for future comparative studies aiming at understanding the evolution, the development, and the function of the larvacean house.

The production, the structure and the function of the filter-feeding house can vary widely between larvacean families^13^, and also between species that belong to the same genus^40^. How these differences could impact animal growth and reproductive success remain largely speculative. For instance, it has often been proposed that feeding in oikopleurids should be less cost-effective, since a fraction of the resources extracted during grazing are spent for the renewal of discarded houses. Generation times measured in our laboratory culture are equivalent for *O. dioica* and *F. borealis*, suggesting that feeding behavior does not have a strong influence on the development speed, at least when animals are raised at comparable densities. Instead, lifecycle length could depend more on animal body size and genome content, and indeed *O. dioica* and *F. borealis* are also similar at the levels. Gaining access to the complete lifecycle of other species with larger genomes^16^ could be critically useful to address this question.

Descriptions of the house-producing epithelium have been reported early in the genus *Oikopleura*^41,42^ and unlike *Fritillaria*^19,43^, knowledge about development benefited from descriptions with modern microscopy techniques. Studies comparing different *Oikopleura* species have shown conserved fields of oikoplastic cells^36^ forming a characteristic pattern at the surface of the trunk. Our results also give clear indications about the presence of distinctive, conserved cell arrangements in the *Fritillaria* epithelium with the most conspicuous one being the dorsal pair of rows of cells, that can also be recognized in the zoological drawings of a few other *Fritillaria* species^44,45^. Our study also showed variations between *F. borealis* and *F. haplostoma*, which we suspect to be related to the production of different structures in the house. While the utility of such character to resolve some uncertainties around the classification of *Fritillaria*^46^ remains to be demonstrated, such lineage-specific arrangements of house-producing cells immediately raises questions about the evolution of the epithelium in larvaceans and the nature of the oikoplastic organ in the common ancestor. The *F. borealis* genome has at least one *CesA* gene closely related to the tunicate homologues (**FigS**)^47^. In contrast, we could not detect any clear homology with oikosins that have been characterized in *O. dioica*^48^, suggesting significant divergence in the complement of structural proteins of the house. Examining the expression of transcription factors involved in the patterning of the trunk epithelium^5^ could bring more informative insight for understanding how this organ evolved in different families of larvaceans.

Although larvaceans embryos can be recovered from seawater samples, producing them from adults that have been isolated and maintained in culture presents several advantages. It allows larger amounts of animals to be examined and it also helps to dissipate uncertainties about the species to which the embryos belong. This is of critical importance since larvae of *Fritillaria* and *Oikopleura* can appear very similar when examined superficially^49^. To our knowledge, our study is the first to provide detailed observations of the *Fritillaria* larva, which revealed developmental processes that would have been unsuspected if the study was based solely on adult specimen. However, without further experimental manipulation and molecular insights, we can only speculate about the responsible mechanisms. For example, even though newly hatched *F. borealis* and *O. dioica* larvae look generally similar, they already show differences in trunk shape with the former having more elongated trunk than the latter. It suggests that the molecular decisions controlling the patterning of the trunk could happen quite early before hatching. Our results also suggest that AP signaling could regulate the development of the epithelial layer. We saw indeed a striking antagonism between the anterior part of pharyngeal trunk epithelium ectoderm, where patterns of house-secreting cells appear early (**Fig6A)**, and the posterior part that undergoes extensive rearrangement before fields of oikoplastic cells are established. Another obvious indication for a role of AP signaling is the progressive, anterograde dissociation of the epithelium that we observed during the development of *F. borealis* larvae. We suppose that AP signaling could regulate the digestion of the basal lamina surrounding the gut and the esophagus, and the dissociation of the epithelial cell layer. Cell death could also participate to the removal of epithelium from these organs. However, since we observed that a high number of cells remained in the posterior pharyngeal trunk epithelium after the anterograde retraction, we suppose that it plays only a minor role. The role of AP signaling for epithelium development would be best addressed using *O. dioica*, where different gene manipulation techniques have been recently established^50^. However, the elongated shape of the *F. borealis* trunk could prove more advantageous for revealing AP patterning with gene expression profiling based on *in situ* hybridization probes. In combination with live imaging and surface labelling of the larva, such approaches could bring definitive answers about the cellular and molecular mechanisms responsible for epithelium development in *Fritillaria*.

## Notes

### Competing Interest Statement

The authors have declared no competing interest.

